## Supplementary figures and images for "Striatin 3 and MAP4K4 cooperate towards oncogenic growth and tissue invasion in medulloblastoma"

### S10, uncropped IB

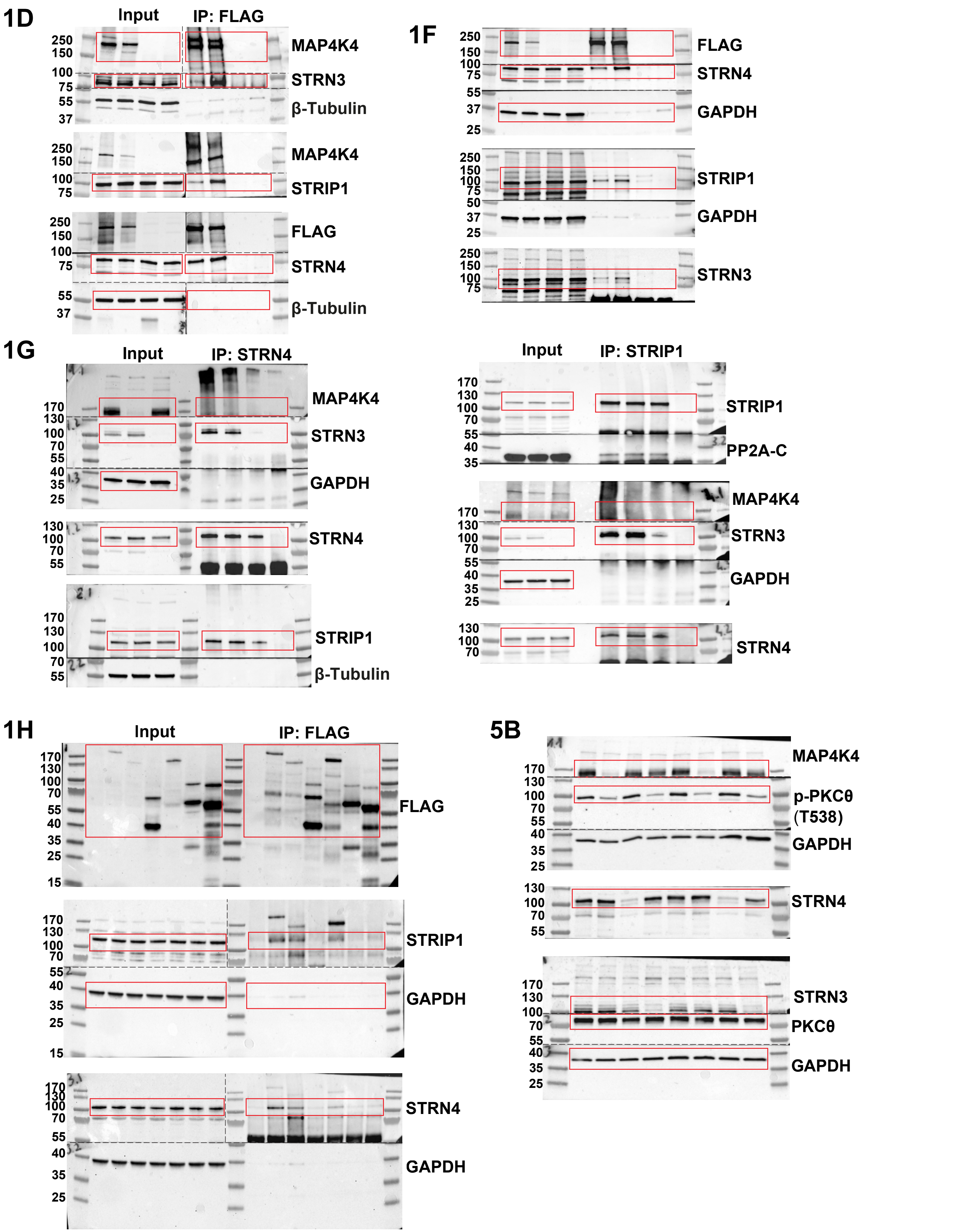

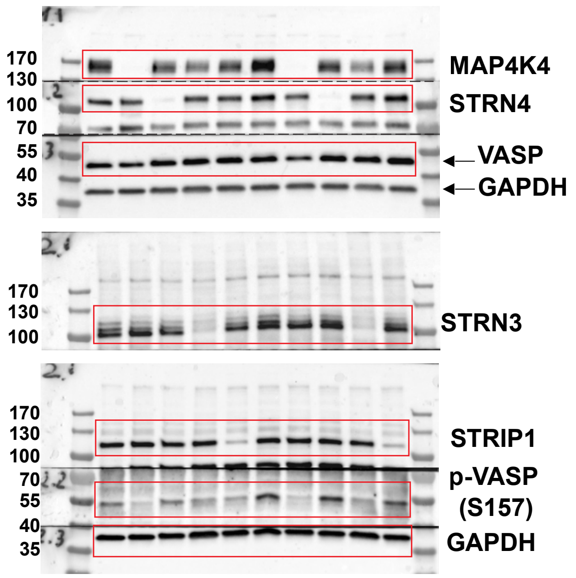

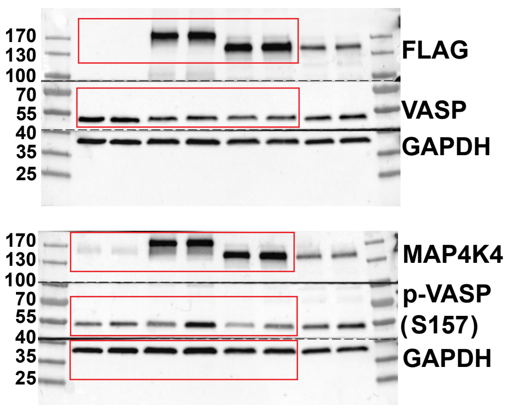

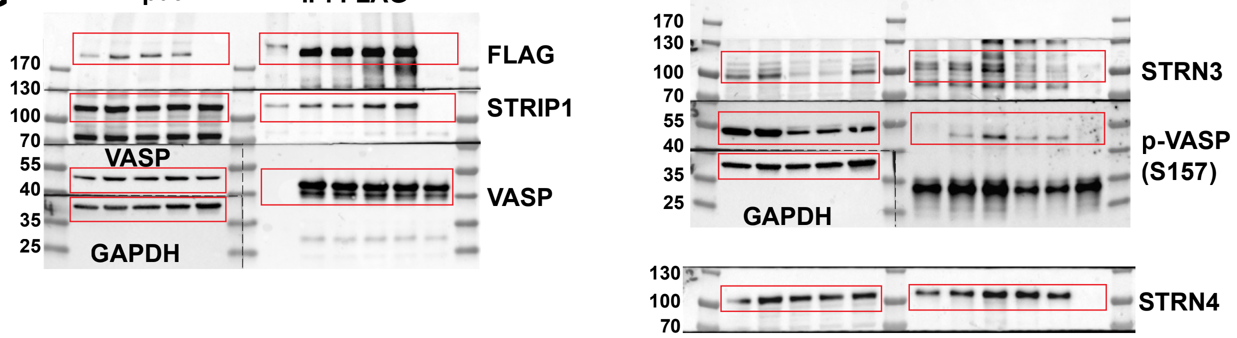

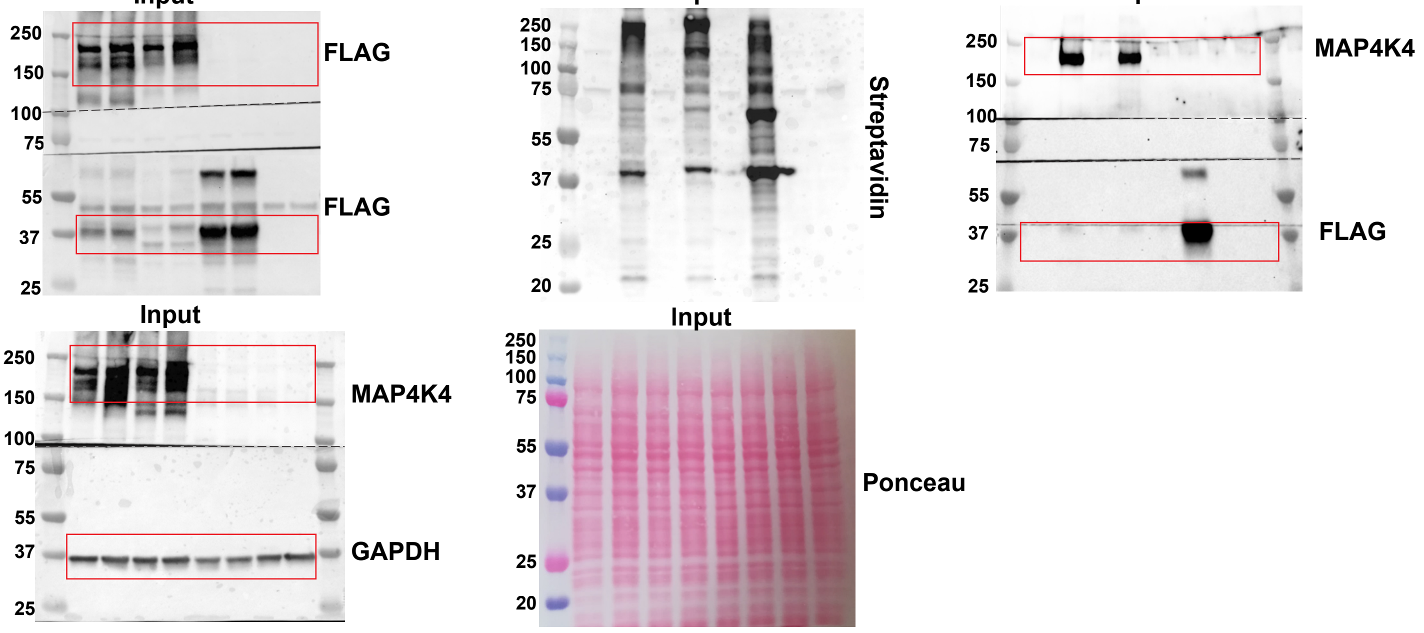

S1D

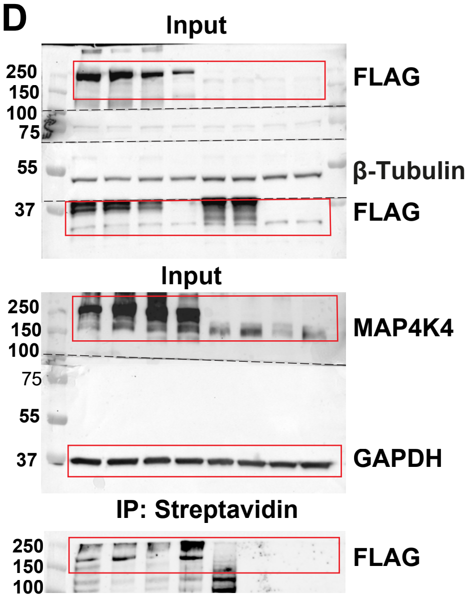


S1G

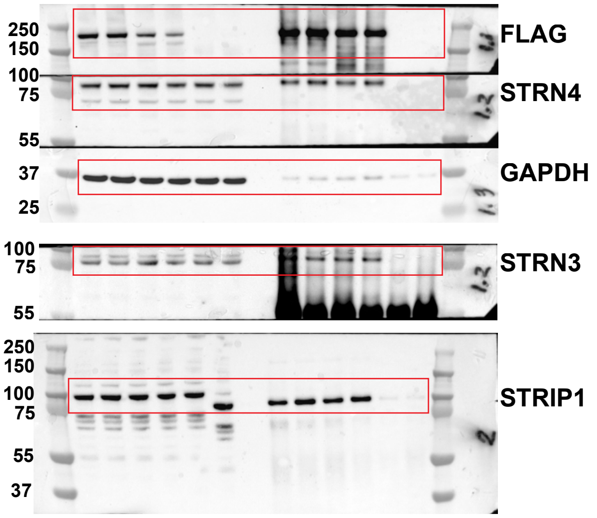


S3A

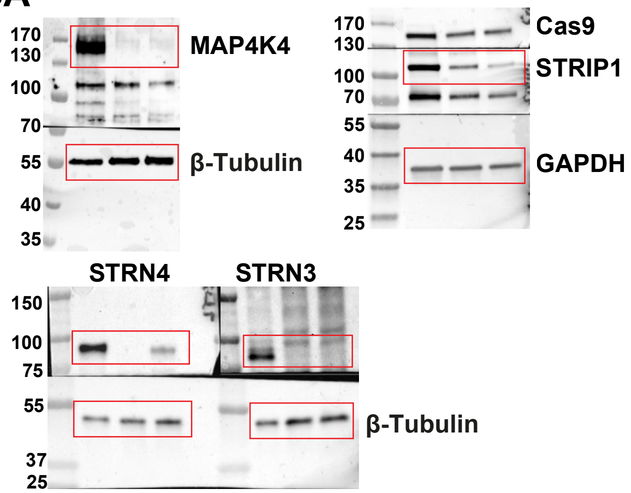


S3B

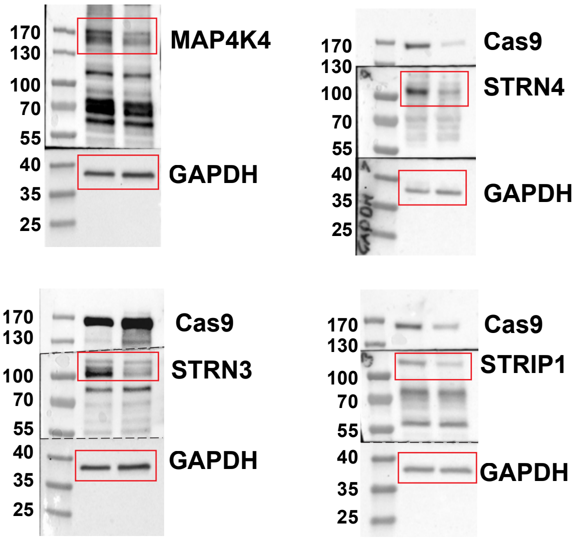


S3E

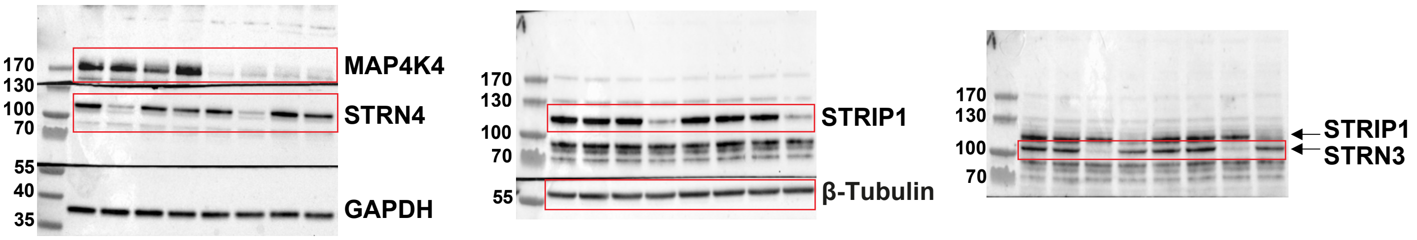


S3G

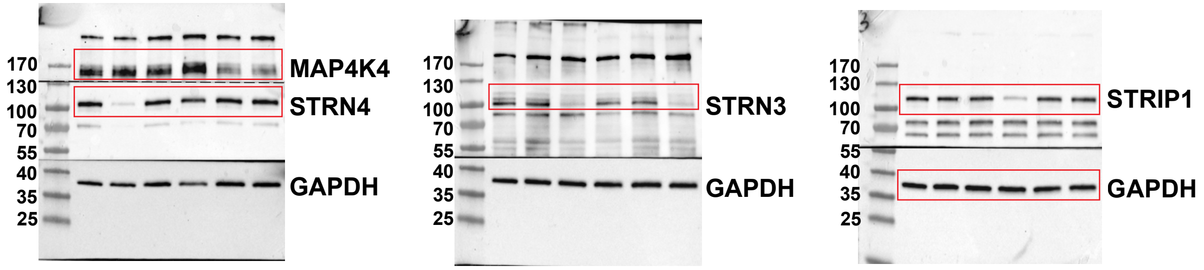

S8H

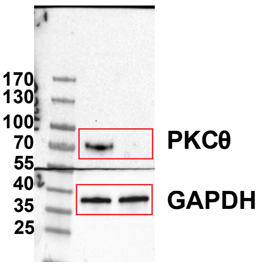


S9D

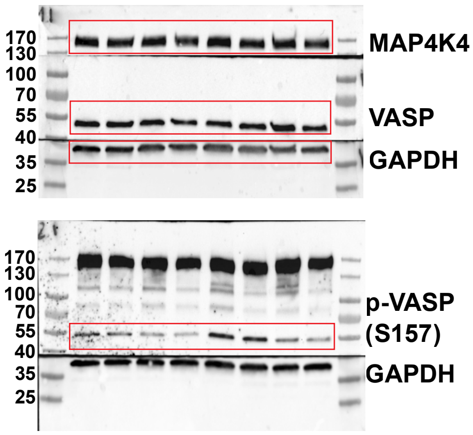


S9E

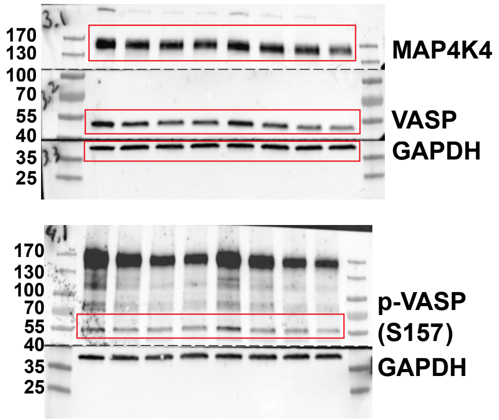


S9G

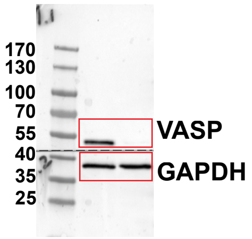
