## Supplementary materials for "Striatin 3 and MAP4K4 cooperate towards oncogenic growth and tissue invasion in medulloblastoma"

**Table 1**: Antibodies used in this study

| **Antibodies** | | | |
| --- | --- | --- | --- |
| **Ab name** | **Company** | **Identifier** | **Dilution** |
| Anti-β-tubulin | Sigma Aldrich | T5201 | 1:1000 (IB) |
| Anti-GAPDH (14C10) | Cell signaling technologies | 2118s | 1:2000 (IB) |
| Anti-FLAG® M2 | Sigma Aldrich | F1804 | 1:1500 (IB) 3 μg / mg lysate (IP) |
| Streptavidin-HRP | Jackson Immunoresearch | 16-030-084 | 1:500 (IB) |
| Anti-MAP4K4 | Abcam | ab80418 | 1:2000 (IB) |
| Anti-STRN4 | Abcam | ab194948 | 1:2000 (IB)  3 μg / mg lysate (IP) |
| Anti-STRN3 (S68) | Thermo Fisher Scientific | MA1-46461 | 1:2000 (IB) |
| Anti-STRIP1 | Abcam | ab199851 | 1:2000 (IB)  3 μg / mg lysate (IP) |
| Anti-PKCθ (E1I7Y) | Cell signaling technologies | 13643 | 1:1000 (IB) |
| Anti-phospho-PKC θ (Thr538) | Cell signaling technologies | 9377 | 1:1000 (IB) |
| Anti-VASP (9A2) | Cell signaling technologies | 3132 | 1:1000 (IB)  1: 400 (IFA) |
| Anti-phospho-VASP (Ser157) | Abcam | ab47268 | 1:750 (IB) |
| Anti-mouse HRP linked | Cell signaling technologies | 7076 | 1:5000 (IB) |
| Anti-rabbit HRP linked | Cell signaling technologies | 7074 | 1:5000 (IB) |
| Normal rabbit IgG | Cell signaling technologies | 2729 | 3 μg / mg lysate (IP) |
| easyBlot anti-mouse HRP | GeneTex | GTX221667-01 | 1:1000 (IB) |
| Anti-BioID2 | Novus | NBP2-59940 | 1:200 (IFA) |
| Streptavidin Alexa Fluor 594 conjugate | Invitrogen | S11227 | 1:500 (IFA) |
| Anti-Calbindin | Abcam | ab108404 | 1:1000 (IFA) |
| Anti-GFAP | Abcam | ab53554 | 1:250 (IFA) |
| Anti-human nuclei (3E1.3) | Millipore | MAB4383 | 1:250 (IFA) |
| Donkey anti-Mouse IgG Secondary Antibody, Alexa Fluor 488 | Thermo Fisher Scientific | A32766 | 1:400 (IFA) |
| Cy3-conjugated Donkey anti-Rabbit IgG | Jackson Immunoresearch | 711-165-152 | 1:250 (IFA) |
| Brilliant Violet 421 Donkey Anti-Goat IgG | Jackson Immunoresearch | 705-675-147 | 1:100 (IFA) |
| Goat anti-Mouse IgG Secondary Antibody, Alexa Fluor 405 | Thermo Fisher Scientific | A31553 | 1:200 (IFA) |
| Hoechst | Sigma Aldrich | B2883 | 1:2000 |

**Table 2**: Sequence information for sgRNAs used in this study

| **sgRNA Oligonucleotides** | | |
| --- | --- | --- |
| **Name** | **Exon** | **Target sequence** |
| sgCTRL | - | GTAGCGAACGTGTCCGGCC |
| sgSTRN4#1 | Exon 1 | GCTCAGGTGGCCTTCCTTCA |
| sgSTRN4#2 | Exon 2 | TTCCTTCAGGGAGAGAGGAA |
| sgSTRN3#1 | Exon 2 | AGGTCAAGAGAACCTGAAGA |
| sgSTRN3#2 | Exon 2 | AGTATGCATTAAAACAAGAA |
| sgSTRIP1#1 | Exon 4 | TGCCAGGGAGAAGAGACTCA |
| sgSTRIP1#2 | Exon 4 | GGATGGCTTGGAAGTCACTG |
| sgMAP4K4#1 | Exon 7 | GGGCGGAGAAATACGTTCAT |
| sgMAP4K4#2 | Exon 4 | CAGGACATGATGACCAACTC |

**Table 3**: siRNA sequences used in this study

| **siRNA** | | | |
| --- | --- | --- | --- |
| **Name** | **Target sequence** | **Company** | **Identifier** |
| siCTRL | UAAGGCUAUGAAGAGAUAC | Dharmacon | D-001210-02-05 |
| siSTRN4 | GGAUCAAGAUGCUAGAGUA | Dharmacon | D-020389-01-0002 |
| siSTRN3 | GGAGGAGGCAAGUCAUUUA | Dharmacon | D-019145-01-0002 |
| siSTRIP1 | GCAGCAAAUUUAUAGGUUA | Dharmacon | D-021516-01-0002 |
| siMAP4K4 | UAAGUUACGUGUCUACUAU | Dharmacon | D-003971-05-0002 |
| siPRKCQ | GCGAGGCUGUUAACCCUUA | Dharmacon | D-003525-05-0002 |
| siVASP | GGGCCACUGUGAUGCUUUA | Dharmacon | D-019763-01-0002 |

**Table 4**: Custom qPCR primers used in this study

| **qPCR primers** | | | |
| --- | --- | --- | --- |
| **Target genes** | | **Sequence** | **Product size** |
| *18S* | F | GGATGTAAAGGATGGAAAATACA | 23 bp |
|  | R | TCCAGGTCTTCACGGAGCTTGTT |  |
| *MAP4K4* | F | GTTACACTAATGCGCACCAC | 193 bp |
|  | R | GTACTTGCCACCAGTCTGCT |  |
| *STRN4* | F | CTCAGGTGGCCTTCCTTCAG | 20 bp |
|  | R | TTTGGCCCTTTCCTGCTTCA |  |
| *STRN3* | F | TGGCACAGAATGGGCTGAAC | 101 bp |
|  | R | CTCCAAGGCCCAGTACACTT |  |
| *STRIP1* | F | CGCAAAGACTCAGAGGGCTA | 109 bp |
|  | R | GCCCTTCCGTGTAGCTGTAA |  |
| *CTGF* | F | CACCCGGGTTACCAATGACA | 119 bp |
|  | R | GGATGCACTTTTTGCCCTTCTTA |  |
| *CYR61* | F | ACAGCAGCCTGAAAAAGGGC | 104 bp |
|  | R | GGGCCGGTATTTCTTCACACT |  |
| *ANKRD1* | F | TAGCGCCCGAGATAAGTTGC | 97 bp |
|  | R | GTCTGCCTCACAGGCGATAA |  |
| *PRKCQ* | F | AACTTTGACTGCGGGTCCTG | 100 bp |
|  | R | TCTGCCCGTTCTCTGATTCG |  |
| *VASP* | F | GGAAGAGATGAACGCCATGC | 94 bp |
|  | R | CTCTGGCTCCTCATTGGCAG |  |

Supplementary figures

**Figure S1: BioID identified MAP4K4 interactome. (A)** Schematic diagram of BioID technology and schematic representation of the lentiviral vectors used to generate 3xFLAG-tagged MAP4K4-BioID2 cell lines. Biotin ligase (BioID2) was fused to either the N-terminus (FLAG-BioID2-MAP4K4) or C-terminus (MAP4K4-BioID2-FLAG) of MAP4K4. An extended flexible linker consisting of 13 repeats of GGGGS was inserted between the coding regions of MAP4K4 and BioID2. As negative control, BioID2-FLAG was used (adapted from^82^). **(B)** Immunoblot analysis (IB) of HEK-293T total cell lysate or after immunoprecipitation (IP) with streptavidin-conjugated beads and detection with HRP-streptavidin. **(C)** Venn diagram of affinity captured proteins from HEK-293T cells. Comparison of BioID2-MAP4K4 with BioID2-CTRL is shown. Values of enriched proteins (p < 0.05, one-way ANOVA) are shown. *n*=3. **(D)** IB of FLAG-tagged BioID2 alone or fused to MAP4K4 in DAOY cell lysate or after pull-down with streptavidin-conjugated beads. **(E)** Confocal microscopy images of DAOY cells expressing BioID2-MAP4K4 fusion protein or BioID2 alone. Biotinylated proteins were labeled with streptavidin (red) and predominantly co-localized with BioID2 (green). Considerable biotinylation is observed in biotin (50 μM) supplemented cells expressing BioID2. DNA is labeled with Hoechst (blue). Scale bar: 30 μm. **(F)** Venn diagram of affinity-captured proteins from DAOY BioID2-MAP4K4 cells compared to BioID2-CTRL. *n*=2. **(G)** DAOY cells expressing BioID2-MAP4K4 fusion protein were serum-starved overnight and treated with 1 µM GNE-495 for 16 h where indicated. FLAG-immunoprecipitated MAP4K4 was subjected to immunoblot analysis for STRIPAK components. GAPDH was used as loading control.

**Figure S2: STRIPAK complex members are highly expressed in MB patients. (A, B)** mRNA (A) and protein (B) levels of STRN4, STRN3, and STRIP1 in a cohort of 218 pediatric brain tumor samples representing seven histological types, including medulloblastoma (MB, *n*=22), low-grade glioma (LGG, *n*=93), high-grade glioma (HGG, *n*=25), ependymoma (EP, *n*=32), craniopharyngioma (CP, *n*=18), ganglioglioma (GG, *n*=18), and atypical teratoid rhabdoid tumor (ATRT, *n*=12). The graphs represent the Z-score values for protein and mRNA in each group. * p < 0.05, ** p < 0.01, *** p < 0.001, **** p < 0.0001 (one-way ANOVA). **(C)** Heatmap for protein or phosphoprotein levels of the indicated proteins in the same cohort of patients as in A. **(D)** Scatterplot showing the phospho-abundance of the indicated residues of MAP4K4 versus the protein expression levels of STRN4, STRN3, or STRIP1 in MB samples. The data were obtained from the CPTAC data portal (<http://pbt.cptac-data-view.org/>).

**Figure S3: CRISPR/Cas9 and siRNA-mediated downregulation of MAP4K4 and STRIPAK complex members. (A, B)** Immunoblots showing CRISPR/Cas9-mediated depletion of MAP4K4, STRN4, STRN3, or STRIP1 in DAOY (A) or HD-MBO3 (B) cells. The numbers indicate the sgRNA used for each gene. **(C)** Invasion and representative images of 3D spheroid invasion assay (SIA) of DAOY cells KO for MAP4K4, STRN4, STRN3, or STRIP1 ± bFGF (100 ng/ml) (*n*=3, means ± SEM). **(D)** qRT-PCR analysis of *MAP4K4*, *STRN4*, *STRN3*, and *STRIP1* in DAOY cells 72 h after transfection with the indicated siRNA (*n*=3, means ± SEM). **(E)** IB analysis and quantification of MAP4K4, STRN4, STRN3, or STRIP1 protein expression level in DAOY cells 72 h after transfection with the indicated siRNA (*n*=4, means ± SEM). **(F)** qRT-PCR analysis of *MAP4K4*, *STRN4*, *STRN3*, and *STRIP1* in UW228 cells 72 h after transfection with the indicated siRNA (*n*=3, means ± SEM). **(G)** Immunoblot analysis and quantification of MAP4K4, STRN4, STRN3, or STRIP1 protein expression level in UW228 cells 72 h after transfection with the indicated siRNA (*n*=3, means ± SEM). **(H)** CellTox Green viability analysis of DAOY spheroids treated for 24, 48, and 72 h with increasing concentrations of LB-100. (*n*=3, means ± SD). Statistical analyses in C, D, F were performed by one-way ANOVA, in E and G by unpaired Student’s t-test. * p < 0.05, ** p < 0.01, *** p < 0.001, **** p < 0.0001.

**Figure S4: Depletion of MAP4K4 and STRN3 blocks tissue invasion in SHH MB cell model. (A)** Timeline of organotypic cerebellum slice culture (OCSC) and tumor spheroid implantation. **(B)** High-resolution (63x) images of OCSCs co-cultured with siRNA-transfected DAOY tumor cell spheroids 72 h after transfection without GF treatment. Green: Lifeact-EGFP; blue: calbindin (Purkinje cells); red: GFAP; yellow: Edu-Click-IT. Scale bar: 50 μm. **(C)** Maximum intensity projections (MIP) of representative confocal sections of OCSCs implanted with DAOY tumor cell spheroids and treated with 30 ng/ml EGF for 48h. Green: Lifeact-EGFP; blue: calbindin (Purkinje cells); red: GFAP. Middle: Inverted greyscale images of MIP of LA-EGFP channel, lower: Inverted greyscale images of MIP of EdU-Click-IT staining. Scale bar: 200 μm. **(D)** Quantification of LA-EGFP area of spheroids shown in C (*n*=3, means ± SD). **(E)** Number of EdU-positive nuclei normalized by the area of the tumor spheroids shown in C (*n*=3, means ± SD). * p < 0.05, ** p < 0.01 (unpaired Student’s *t*-test).

**Figure S5: High-resolution images of organotypic cerebellum slice culture implanted with HD-MBO3 spheroids.** Single confocal microscopy sections of OCSCs implanted with tumor spheroids derived from HD-MBO3 with CRISPR/Cas9-mediated knockout of MAP4K4, STRN3, or STRIP1. 63x image acquisition of slices five days after implantation and treatment with 12.5 ng/ml bFGF. 4x magnifications of boxed areas are shown. Green: Lifeact-EGFP; blue: calbindin (Purkinje cells); red: human nuclei; yellow: Edu-Click-IT. Scale bar: 200 μm.

**Figure S6: STRIPAK enables FGFR-PP2A-mediated activation of YAP/TAZ transcriptional program. (A)** Models for MAP4K4 and STRIPAK regulation of Hippo pathway and YAP/TAZ target gene expression. **(B)** qRT-PCR analysis of *CYR61* (upper) and *ANKRD1* (lower) expression in DAOY cells 48 h after transfection with the indicated siRNA ± treatment for 4 h with 5 µM LB-100 (*n*=3, means ± SEM). **(C, D)** qRT-PCR analysis of *CYR61* and *ANKRD1* expression in DAOY (C) or HD-MBO3 cells (D) transfected with siRNA for 48 h, serum-starved overnight, and treated with 100 ng/ml bFGF for 1 h and/or 5 µM LB-100 for 4 h where indicated (*n*=3, means ± SEM). * p < 0.05, ** p < 0.01, *** p < 0.001, **** p < 0.0001 (unpaired Student’s *t*-test).

**Figure S7:** **Phylogenetic distribution of the kinases predicted the peptide chip array**. **(A)** Upstream kinase prediction analysis of DAOY cells transfected with the indicated siRNA ± treatment with 100 ng/ml bFGF. The plots show the top differentially activated putative upstream tyrosine kinases (PTK) predicted to phosphorylate the phosphosites on the PamChip®. The x-axis indicates the values for the mean kinase statistic, which represents the difference in the activity of the predicted protein kinase between the two compared groups, with effect size (values) and direction (>0=activation; <0=inhibition). The color of the bars represents the specificity score (darker the color, higher the specificity). Values of the specificity score >0.9 were considered statistically relevant. First graph from left: comparison of bFGF stimulated siCTRL cells vs. untreated (UT); all other graphs: comparison of the indicated siTarget vs. siCTRL in the presence of bFGF. *n*=3. **(B-E)** Phylogenetic kinome trees illustrating the family distribution of the upstream kinases predicted to phosphorylate the serine/threonine (STK) and protein tyrosine (PTK) consensus peptides of the PamChip®. Colored dots highlight the kinases where predicted activity is significantly different in the indicated siTarget (B: siMAP4K4; C: siSTRN4; D: siSTRN3, E: siSTRIP1) compared to siCTRL in bFGF stimulated conditions in DAOY cells. The coloring scale is based on the mean kinase statistic and ranges from −2 (strong decrease of kinase activity in siTarget vs. siCTRL, red color) to +2 (strong increase of kinase activity in siTarget vs. siCTRL, green color). The size of the circle represents the specificity score.

**Figure S8: Novel protein kinases C are pro-migratory effectors of MAP4K4 and STRN3. (A-C)** Quantification and representative images of SIA with DAOY cells stimulated with 100 ng/ml bFGF and treated with increasing concentrations of Darovasertib (PKCθ inhibitor, A), Rottlerin (PKCδ inhibitor, B) or Bisindolylmaleimide I (BIM, PKCα/β/γ inhibitor, C). Invasion was quantified as the sum of invasion distances from the center of the spheroids. **(***n*=4, means ± SEM). **(D)** Maximum intensity projections (MIP) of representative confocal sections of OCSCs implanted with HD-MBO3 tumor cell spheroids and treated with 5 µM PKCθ inhibitor, 5 µM Rottlerin, or DMSO for five days. Green: Lifeact-EGFP; blue: calbindin (Purkinje cells); red: GFAP; yellow: Edu-Click-IT. Scale bar: 200 μm. **(E)** Quantification of LA-EGFP area of HD-MBO3 spheroids shown in D (*n*=3, means ± SD). **(F)** Quantification of the number of EdU-positive nuclei normalized by the area of HD-MBO3 spheroids shown in D (*n*=3, means ± SD). **(G)** qRT-PCR analysis of *PRKCQ* (PKCθ) mRNA expression in DAOY cells 72 h after siRNA transfection (*n*=4, means ± SEM). **(H)** IB analysis and quantification of siRNA-mediated PKCθ depletion in DAOY cells 72 h after transfection (*n*=3, means ± SEM). **(I)** MIP of representative confocal sections of OCSC implanted with transfected HD-MBO3 tumor cell spheroids and treated with 12.5 ng/ml bFGF for 48h. Green: Lifeact-EGFP; blue: calbindin (Purkinje cells); red: GFAP. Scale bar: 200 μm. **(J)** Quantification of LA-EGFP area of HD-MBO3 spheroids shown in I (*n*=3, means ± SD). Statistical analyses in A-C were performed by one-way ANOVA, in E-H and J by unpaired Student’s t-test. * p < 0.05, ** p < 0.01, **** p < 0.0001.

**Figure S9: MAP4K4 mediates VASP_S157_ phosphorylation. (A, B)** Individual volcano plots representing the changes in phosphorylation of phospho-serine/threonine (STK, A) and phospho-tyrosine (PTK, B) peptides in the indicated siTarget versus siCTRL in bFGF-stimulated DAOY cells. The p-values were calculated versus siCTRL by ANOVA and post-hoc Dunnett’s test in the BioNavigator software. *n*=3. The peptides that showed significant changes in phosphorylation are highlighted in red (p < 0.05) or orange (p < 0.1). **(C)** Maximum intensity projection of LA-EGFP fluorescence of tissue-invading HD-MBO3 CTRL or KO MAP4K4 or STRN3 cells. Scale bar: 10 µm. Arrowheads indicate filopodia-like protrusions. **(D-E)** IB and quantification of VASP_S157_ phosphorylation in DAOY (D) and HD-MBO3 cells (E) after treatment for 4 h with increasing concentration of GNE-495 ± 100 ng/ml bFGF for 15 min. Quantification represents the ratio of p-VASP_S157_ over total VASP relative to untreated control (B: *n*=3, means ± SEM. C: *n*=2, means ± SEM). **(F)** qRT-PCR analysis of *VASP* mRNA expression in DAOY cells 72 h after siRNA transfection (*n*=4, means ± SEM). **(G)** IB analysis and quantification of siRNA-mediated VASP depletion in DAOY cells 72 h after transfection (*n*=3, means ± SEM). Statistical analysis in D and E was performed by one-way ANOVA, in F and G by unpaired Student’s t-test. * p < 0.05, ** p < 0.01, **** p < 0.0001.

**Figure S10: Original uncropped immunoblot images**
